# The salivary virome during childhood dental caries

**DOI:** 10.1101/2024.05.22.595360

**Authors:** Jonah Tang, Jonathon L. Baker

## Abstract

While many studies have examined the bacterial taxa associated with dental caries, the most common chronic infectious disease globally, little is known about the caries-associated virome. In this study, the salivary viromes of 21 children with severe caries (>2 dentin lesions) and 23 children with healthy dentition were examined. 2,485 viral metagenome-assembled genomes (vMAGs) were identified, binned, and quantified from the metagenomic assemblies. These vMAGs were mostly phage, and represented 1,547 unique species-level vOTUs, 247 of which appear to be novel. The metagenomes were also queried for all 3,835 unique species-level vOTUs of DNA viruses with a human host on NCBI Virus, however all but *Human betaherpesvirus 7* were at very low abundance in the saliva. The oral viromes of the children with caries exhibited significantly different beta diversity compared to the oral virome of the children with healthy dentition; several vOTUs predicted to infect *Pauljensenia* and *Neisseria* were strongly correlated with health, and two vOTUs predicted to infect Saccharibacteria and *Prevotella histicola*, respectively, were correlated with caries. Co-occurrence analysis indicated that phage typically co-occurred with both their predicted hosts and with bacteria that were themselves associated with the same disease status. Overall, this study provided the sequences of 53 complete or nearly complete novel oral phages and illustrated the significance of the oral virome in the context of dental caries, which has been largely overlooked. This work represents an important step towards the identification and study of phage therapy candidates which treat or prevent caries pathogenesis.

**Importance:** Dental caries is the most common chronic infectious disease, worldwide, and is caused by a dysbiosis of the oral microbiome featuring an increased abundance of acid-tolerant, acid-producing, and biofilm-forming bacteria. The oral microbiome also contains viruses; however, very little is known about the the caries-associated virome. In this study, the salivary virome of children with severe caries was compared to the salivary virome of children with healthy dentition. The metagenomes contained a total of 1,547 unique species-level vOTUs, 247 of which appeared to be novel. The viromes from the children with caries were significantly different than the viromes from the children with healthy teeth, and several health- and disease-associated vOTUs were identified. This study illustrated the importance of the oral virome in the context of dental caries, and serves as a step towards a better understanding of oral inter-kingdom interactions and identification of potential phage-based caries therapeutics.

## Main Text

The human oral microbiome is comprised of bacteria, fungi, viruses (including phages), archaea, and microeukaryotes, and has a major impact on health, with dental caries, periodontal disease, and oral (head and neck) cancers having mainly microbial etiologies (1). Dental caries, in particular, is the most common chronic infectious disease, worldwide, and is caused by a dysbiosis of the dental plaque microbiome characterized by an abundance of bacterial taxa that are excellent biofilm formers with an ability to generate and then tolerate high concentrations of acids which destroy the tooth enamel (with the archetype taxa being *Streptococcus mutans*) (2). In addition to its role in oral health and disease, the oral microbiome is increasingly recognized as playing an important role in systemic/overall health as well (3). The overwhelming majority of oral microbiome studies, and microbiome studies overall, have focused on the bacterial constituents. In many cases this is due to analysis based on 16S rRNA amplicon sequencing, which ignores the non-bacterial members of the microbiome. Furthermore, when metagenomics approaches have been employed, they have been mainly dependent on either read-based analyses using databases focused on known bacterial taxa or on *de novo* assembly-based approaches dependent on metagenome-assembled genome (MAG) binning and quality control tools also optimized for bacteria. As a result, study of the virome component of the human microbiome, and its impact, has lagged behind, despite early landmark studies illustrating that both bacteriophage and human viruses contribute to the health of the oral microbial community and the overall health of the human host (4–6). A previous study by our group used deep metagenomics to examine the oral microbiome in advanced childhood caries compared to health (7). In that study, taxonomic abundance analysis using MetaPhlaAn2 (8) provided some rudimentary insight into the oral virome, and even suggested that *Human gammaherpesvirus 4* (Epstein-Barr Virus [EBV]) was associated with dental caries. However, a specific virus-oriented analysis was not performed, and only bacterial genomes were analyzed from the *de novo* metagenomic assemblies (due to a lack of virus-oriented tools at the time).

In this study, we reanalyzed 44 assembled oral metagenomes (21 from children with advanced caries and 23 from children with healthy dentition) using the recently developed ViWrap pipeline, which uses genome identification, binning, and quality analysis methods optimized for obtaining viral metagenome-assembled genomes (vMAGs) (9) (detailed methods are available in Supplemental Text S1, and all scripts and code associated with this project are available at https://github.com/jonbakerlab/caries-associated-virome). ViWrap identified 2,485 genomes vMAGs across these assemblies (Figure 1, Supplemental Table S1). The vast majority of these vMAGs were viral taxa within the Caudovirales class (Figure 1). Several notable exceptions were 8 Microviridae, 1 Adenoviridae, 2 Mimiviridae, 1 Pokkesviricetes, and 1 *Human betaherpesvirus* vMAGs. The 2,485 vMAGs were dereplicated to remove redundancy at ≥95% average nucleotide identity (ANI), resulting in 1,547 species-level viral genome clusters (vOTUs; Figure 1, Supplemental Table S2). To determine the novelty of these genomic sequences, the 1,547 vOTUs were compared to all ‘Bacteriophages’ from NCBI Virus (10) (42,801 genomes, as of April 2024), as well as the recently developed Oral Virus Database (OVD; containing 48,425 nonredundant virus genomes) (11). 247 of the nonredundant vOTUs obtained in this study contained zero hits at an average nucleotide identity (ANI) of ≥95%, indicating that they were novel species-level vOTUs (Figure 1; Supplementary Table S3). Auxiliary metabolic genes (AMGs) are viral genes that augment host functions to benefit the viral infection and/or replication processes. Across the vMAGs, there were 115 predicted AMGs, with 11 AMGs identified uniquely in caries-associated microbiomes and 50 AMGs identified uniquely in health-associated microbiomes (Supplemental Table S4).

**Figure 1:**
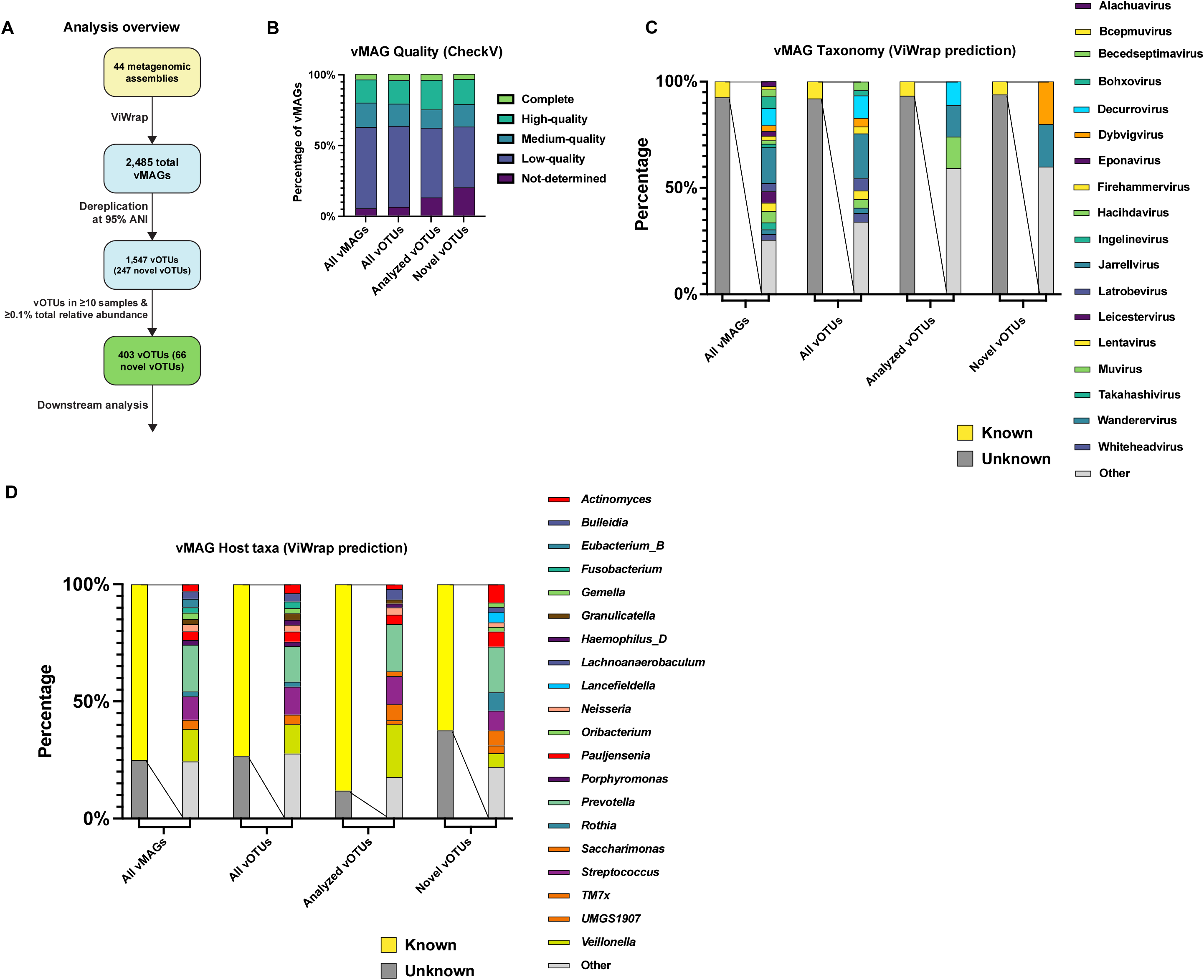
ViWrap analysis of 44 oral metagenomes results in 2,485 vMAGs and 1,547 unique vOTUs, with 247 representing vOTUs not previously described. **(A)** Overview of the approach and viral genomes obtained. 44 oral metagenome assemblies were analyzed with the ViWrap pipeline, which identifies and annotates viral sequences, performs binning and quality analysis, and predicts viral and host taxonomy. This resulted in 2,485 vMAGs, which were dereplicated at ≥95% ANI using Anvi’o, resulting in 1,547 species-level vOTUs. These vOTUs were compared with all bacteriophage genomes on NCBI Virus and in the OVD database using skani, which illustrated that at ≥95% ANI, 247 of the vOTUs did not have representatives in NCBI or OVD, and are therefore likely to be novel. Of the vOTUs obtained, 403 were present in ≥10 samples and were at ≥0.1% total relative abundance across all samples, these were subjected to the downstream analyses. **(B)** vMAG Quality. Bar graph illustrating the quality level of all vMAGs, all vOTUs, analyzed OTUs, and novel OTUs, as determined by CheckV. **(C)** Predicted vMAG taxonomy. Bar graphs illustrating the viral taxonomy, predicted by ViWrap, of all vMAGs, all vOTUs, analyzed vOTUs, and novel vOTUs. **(D)** Predicted host taxonomy. Bar graphs illustrating the taxonomy of the bacterial host, predicted by ViWrap, of all vMAGs, all vOTUs, analyzed vOTUs, and novel vOTUs.

Because ViWrap was optimized to identify phage, and the previous MetaPhlAn2 analysis had also observed human DNA virus genomes in the oral microbiomes (7), we also sought to quantify the abundance of known human viruses in our metagenomes. The 12,482 genomes of DNA viruses with a human host on NCBI Virus (10) (as of March 2024) were also dereplicated to ≥95% ANI, resulting in 3,835 vOTUs (Supplementary Table S5). The metagenomic reads from the 44 samples were mapped to these human vOTUs, which illustrated that most were of very low abundance across our dataset (Supplementary Table S6). 19 of the human vOTUs had ≥10,000 total reads mapped across all samples, and these features were added to the *de novo* assembled vOTUs for 1,566 total vOTU features for downstream analysis. To determine the relative abundances of these vOTUs across the metagenomic samples, while normalizing for genome size, the metagenomic reads were mapped to the 1,566 viral genomes using CoverM (https://github.com/wwood/CoverM) (Supplemental Table S6). 404 vOTUs with a total relative abundance ≥0.1% across all samples and present in ≥10 samples were used for downstream analysis (Supplemental Table S6). *Human betaherpesvirus 7* was the only human DNA virus from NCBI Virus passing these cutoffs, present in 35 samples and at 0.13% total abundance across all samples.

Beta diversity analysis indicated a significant difference between the viromes of the subjects with caries versus the subjects with healthy teeth (PERMANOVA p-value: 0.008; Figure 2A). The only sample metadata with a significant impact on alpha diversity of the viromes was the concentration of salivary epidermal growth factor (p-value: 0.0384). Due to the issues inherent with differential abundance analysis of compositional samples (12), we used two different approaches: differentials calculated by machine learning via Songbird (13) (Supplemental Table S7), and negative binomial distributions via DESeq2 (14) (Supplemental Table S7). Both methods identified 7 vOTUs that had increased abundance correlated with healthy dentition, with two predicted to infect *Pauljensenia*, two predicted to infect *Neisseria*, and 3 with unknown hosts (Figure 2B). Only two vOTUs were significantly increased in caries, and their Songbird ranks were more modest than the vOTUs identified by both methods as increased in health, with one of these caries-associated vOTUs infecting Saccharibacteria and the other infecting *Prevotella histicola* (Figure 2B). MMvec (15) was utilized to examine the co-occurrences of the viral features with both the host immunological markers (Figure 2C, Supplemental Table S8) and taxonomic features (nearly all bacterial; Figure 2D and 2E, Supplemental Table S8) from the previous study (7). Overall, vOTUs tended to co-occur with their predicted bacterial hosts (Figure 2D). vOTUs also tended to co-occur with bacterial OTUs with a similar Songbird differential ranking with respect to being caries-associated or health-associated (i.e. caries-associated vOTUs and caries-associated bacteria largely occurred together; Figure 2E).

**Figure 2:**
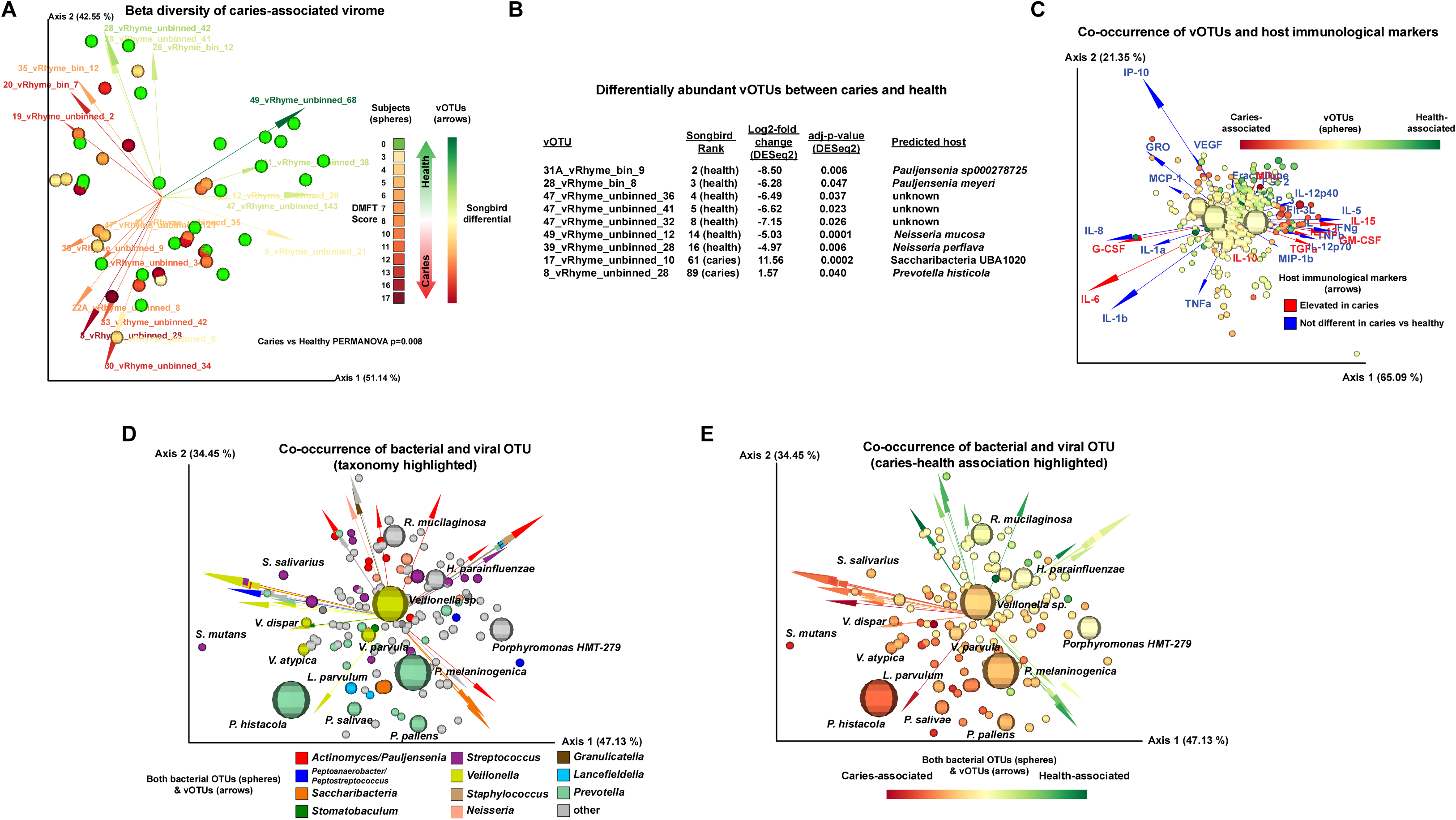
Significant differences in the oral virome, and its co-occurrences, between healthy children and children with caries. **(A)** Beta diversity. Biplot generated by DEICODE (robust Aitchison PCA) illustrating subjects represented as spheres, which are colored with a gradient indicating DMFT score (i.e., caries severity). Feature loadings (i.e., vOTUs driving the differences in the graph ordination space) are illustrated by the vectors, which are color coded based on their Songbird ranks indicating association with caries versus health. **(B)** vOTUs most correlated with health or caries. Table listing the 9 vOTUs that had relative abundance significantly correlated with either health or caries. ‘Songbird Rank’ indicates rank of differential from the extreme (i.e., “61 (caries)” indicates the taxa was the 61^st^ in terms of differential associated with caries). **(C)** Viral-immune marker co-occurrence. Biplot illustrating the co-occurrence of vOTUs with immune markers. vOTUs are represented as spheres, with size indicating total abundance of the taxa across all samples, and color representing Songbird ranks indicating association of caries versus health. Vectors represent host immune markers, with red vectors indicating host immune markers that were significantly elevated in caries, and blue vectors indicating immune markers that were not significantly different between caries and health, as described in (7). **(D and E)** Oral virome-bacteriome co-occurrences. Biplots illustrating the co-occurrence of oral vOTUs with the oral metagenomes (e.g., nearly all bacteria) identified and analyzed in (7). Bacterial taxa are represented as spheres, with size indicating the total abundance of the taxa across all samples. vOTUs are represented by the vectors. In **(D)**, colors of the spheres indicate bacterial taxonomy while colors of the vectors indicate the predicted host taxa of the vOTU (e.g., *Veillonella* bacterial OTUs are yellow spheres and yellow arrows are vOTUs predicted to infect *Veillonella*. In **(E)**, colors of both the spheres and vectors illustrate Songbird ranking indicating association with either caries or good dental health (e.g., red spheres are bacterial taxa highly associated with caries and red arrows are vOTUs highly associated with caries). Interactive QZV files enabling readers to examine the datasets from Figure 2, Panels A, C, D, and E in 3-D, visualize metadata in different ways, and click on individual data points for more information are available at https://github.com/jonbakerlab/caries-associated-virome.

Altogether, this study illustrated the importance of the virome component of the microbiome in the context of dental caries, and the concept that there is still a wealth of unknown viral diversity within the oral microbiome. Limitations of this study include the relatively small and homogenous population size, and the fact that many of the vOTUs obtained were incomplete fragments. Continued improvements in long-read sequencing technology concurrent with a continued reduction in sequencing costs will enable larger-scale future metagenomics studies producing more MAGs with greater contiguity. These types of future studies will not only enable an improved understanding of the dynamics of the oral virome and overall microbiome during caries, but will also hopefully identify promising phages for therapeutic development to treat or prevent this costly disease.

## Data Availability

The raw sequencing reads of the oral metagenomes were originally described in (16) and are available on NCBI with accession numbers PRJNA478018 and SRP151559. The bacterial genomes assembled from those reads were originally described in (7) and are available on NCBI under the accession number PRJNA624185. The 53 vMAGs representing novel vOTUs rated as either ‘complete’ or ‘high-quality’ by checkV are currently at available at https://github.com/jonbakerlab/caries-associated-virome, and are in submission at NCBI GenBank with the accession number to be added here in the final version of this manuscript. All vMAG sequences, vOTU sequences, scripts, code, and interactive QZV files (enabling readers to examine the datasets from Figure 2, Panels A, C, D, and E in 3-D, visualize metadata in different ways, and click on individual data points for more information) are available at https://github.com/jonbakerlab/caries-associated-virome.

## Acknowledgements

This research was supported by NIH/NIDCR R00-DE029228 and by Oregon Health & Science University. A perl script used in the OTU table construction of the human DNA viruses was written by R. A. Richter.

